# Single-nucleus long-read RNA sequencing reveals isoform usage shifts in mouse acute kidney injury

**DOI:** 10.64898/2026.11.10.765759

**Authors:** Yasuhiro Oda, Megan L. Noonan, Eryn E. Dixon, Haojia Wu, Benjamin D. Humphreys

**Affiliations:** Division of Nephrology, Department of Medicine, Washington University in St. Louis, St. Louis, MO, United States

**Author notes:** Correspondence should be addressed to Benjamin D. Humphreys.

**Keywords:** alternative splicing, acute kidney injury, transcriptomics

## Abstract

**Introduction:** Single-cell and single-nucleus short-read RNA sequencing has elucidated transcriptomic changes across kidney cell types at the gene expression level. However, its limited read length precludes reliable isoform identification, hindering the characterization of individual protein-coding isoforms and non-protein-coding isoforms. We combined long-read sequencing with single-cell and single-nucleus RNA sequencing technologies to investigate isoform regulation in healthy and acutely injured mouse kidneys.

**Methods:** Single-cell and single-nucleus long-read RNA sequencing datasets were generated from kidneys of three healthy adult male C57BL/6J mice. An additional three adult male C57BL/6J mice underwent bilateral renal ischemia-reperfusion injury, and kidneys were subjected to single-nucleus long-read RNA sequencing.

**Results:** A total of 1.3 billion reads were generated, with a median read length of 757–884 bases across samples. Single-nucleus libraries yielded a smaller proportion of reads assigned to isoforms than single-cell libraries, reflecting their larger proportion of intronic reads. Nevertheless, single-nucleus libraries detected more distinct isoforms and greater isoform expression diversity, partly owing to the near absence of mitochondrial gene–derived transcripts. Single-nucleus analysis of healthy and injured kidneys revealed that ischemia-reperfusion injury downregulated oxidative phosphorylation– associated genes in proximal tubule cells both at the gene expression level and through shifts in isoform composition. Beyond oxidative phosphorylation–associated genes, *Fut9* and *Prrc2b* were identified as genes exhibiting prominent shifts in isoform usage after ischemia-reperfusion injury.

**Conclusion:** Single-nucleus long-read RNA sequencing detected injury-induced shifts in isoform usage in mouse kidney, adding a layer of transcriptomic information that is largely inaccessible to conventional gene-level short-read analysis.

## Introduction

Single-cell short-read RNA sequencing (RNA-seq) has elucidated transcriptomic changes across kidney cell types, providing deep insights into the underlying mechanisms of diverse kidney diseases.^1–7^ However, standard single-cell analyses quantify expression at the gene level, aggregating all gene transcripts, including isoforms with different or no protein-coding potential. Moreover, widely used droplet-based platforms sequence only a short segment near the 3′ or 5′ end of each transcript, so most internal splicing differences cannot be detected. This poses a challenge for investigating alternative splicing and functionally distinct protein isoforms in the kidney.^8–10^ Even full-length short-read approaches infer isoforms indirectly from individual splice junctions, limiting reliable reconstruction of full-length isoforms.^11^

Over the past decade, long-read sequencing has emerged as a technology capable of sequencing much longer RNA and DNA molecules than short-read sequencing. Whereas short-read sequencing typically reads up to 150 consecutive bases (maximum of 300), long-read sequencing can read thousands of consecutive bases, enabling sequencing of full-length RNA transcripts.^11^ This makes it possible to resolve the complete exon structure of individual transcripts and thereby predict whether a transcript is protein-coding and, for genes with multiple protein-coding isoforms, which protein isoform it encodes. Recent studies have applied long-read RNA-seq with single-cell and single-nucleus technologies to quantify alternatively spliced isoforms across cell types in the brain, heart and retina.^12–15^ In the kidney, long-read sequencing has been used to catalog isoforms in microdissected human glomerular and tubulointerstitial compartments.^16^

Herein, we combined long-read RNA-seq with single-cell and single-nucleus technologies to investigate isoform regulation in healthy and acutely injured mouse kidneys. We hypothesized that renal ischemia-reperfusion injury (IRI) induces cell type–specific isoform usage shifts that are not detected by gene-level analysis and that are predicted to alter the encoded proteins.

## Methods

### Animal experiment

Kidneys were collected from 8 to 11-week-old C57BL/6J male mice that underwent either a bilateral renal IRI surgery two days before sacrifice or no surgical intervention. Bilateral IRI was performed as follows. Mice were anesthetized with isoflurane and received a subcutaneous injection of buprenorphine SR for pain control. Body temperature was maintained between 36.5 and 37.5°C throughout the surgery. Flank incisions were made bilaterally, kidney were exposed, and renal pedicle were clamped with nontraumatic microaneurysm clamps (Roboz, Rockville, MD) for 22 minutes. The clamps were removed, the peritoneal layer was sutured, and the flank incisions were closed with wound clips. All animal experiments were approved by the Institutional Animal Care and Use Committee at Washington University in St. Louis (protocol #23-0460). C57BL/6J mice were purchased from the Jackson Laboratory (Bar Harbor, ME).

### Single cell and single nucleus isolation

Methods and associated references^1,17^ are provided in the Supplemental Material.

### cDNA library preparation and long-read RNA-seq

Single-cell and single-nucleus cDNA samples were generated with Chromium Next GEM Single Cell 3′ Reagent Kits v3.1 (10x Genomics, Pleasanton, CA). To generate full-length cDNA, we followed the kit instructions for Gel Beads-in-emulsion generation and cDNA amplification with the following two modifications: the reverse transcription of Gel Beads-in-emulsion at 53°C was extended from the original 45 minutes to 2 hours, and the extension step at 72°C during cDNA amplification was extended from the original 1 minute to 3 minutes. Fragmentation and 10x library construction were not performed. Libraries were prepared from cDNA samples using the Ligation Sequencing Kit V14 (SQK-LSK114, Oxford Nanopore Technologies, Oxford, UK) and sequenced on a PromethION24 instrument on R10.4.1 flow cells (Oxford Nanopore Technologies). A single library was loaded per flow cell. The sequencing was run for at least 72 hours and stopped after the output dropped off. Maximum run time was 90 hours. Key sample metrics are summarized in Supplemental Table 1.

### Expression matrices generation

Raw sequencing data in POD5 files were basecalled with Dorado v.0.9.0 and model dna_r10.4.1_e8.2_400bps_sup@v5.0.0 (Oxford Nanopore Technologies). FASTQ files were processed with EPI2ME wf-single-cell workflow (Oxford Nanopore Technologies) for processes including adapter identification, cell barcode identification and read assignment. Generated expression matrices were imported to R system for statistical computing (version 4.5.1, R Foundation for Statistical Computing, Vienna, Austria) with Seurat v5^18^ for downstream analyses. Transcriptome reference used for the read assignment was the 2024-A mouse GRCm39 (GENCODE vM33/Ensembl110 annotations) dataset built by 10x Genomics. To correspond this, we used the comprehensive gene annotation on the primary assembly from the mouse GENCODE release M33 (GRCm39) as the reference GTF file in the downstream analyses with R. IsoQuant^19^ was used for quantifying exonic and intronic reads.

### Quality control

Gene expression counts and isoform expression counts were stored in two different assays in a single Seurat object. After ambient RNA removal with SoupX v1.6.2^20^ in each assay, barcode rank plots (known as “knee plots”) were generated for unique molecular identifier (UMI) counts of the gene assay, since EPI2ME wf-single-cell workflow did not fully remove empty droplets in some datasets. Droplets with UMI counts below the bottom of the cliff were obvious empty droplets and removed from the datasets (Supplemental Figure 1, Supplemental Table 1). Following cells and nuclei were regarded as low-quality cells/nuclei and removed from the datasets in this order: 1) number of detected genes ≤ 200, 2) mitochondrial transcript percentage ≥ 1% in single-nucleus datasets only, and 3) either the number of detected genes or the number of UMI is in the upper 1 percentile of the remaining cells (Supplemental Table 1).

### Integration of datasets

Three single-cell datasets and three single-nucleus datasets from the same healthy mice were integrated to analyze the difference derived from the two methods. Three single-nucleus datasets from healthy mice and those from mice post-IRI were integrated to analyze transcriptomic changes caused by the intervention. Datasets from individual samples were integrated using the IntegrateLayers function in the Seurat package with the HarmonyIntegration method, followed by unsupervised clustering. Doublets were estimated by DoubletFinder v2.0.6.^21^ All estimated doublets and clusters enriched for estimated doublets were removed from the datasets. Datasets were reintegrated, reclustered and manually annotated for cell types. In the integrated dataset of single-cell and single-nucleus datasets from healthy mice, one low-quality cluster was identified based on low gene counts and the co-expression of marker genes from multiple cell types (Supplemental Figure 2). This low-quality cluster was removed from the dataset for analysis.

### Bioinformatic analysis of proximal tubule cell transcriptome

Differential isoform usage was analyzed with DTUrtle v1.0.2.^22^ Over-representation analysis and gene set enrichment analysis were performed for Kyoto Encyclopedia of Genes and Genomes (KEGG) pathways using enrichKEGG and gseKEGG functions in clusterProfiler v4.16.0.^23^ Average log2 fold change of the expression level was used for ranking genes in gene set enrichment analysis. When FeaturePlot function was used for visualizing the expression of genes and isoforms with low expression levels, cells were plotted in the order of the expression level by using the “order = TRUE” option.

### Isoform expression diversity analysis

The levels of isoform expression diversity per cell and nucleus were calculated as follows.

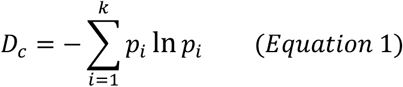

Equation 1 defines the Shannon entropy of a given cell or nucleus (*Dc*), where *k* is the number of splice variants detected in the cell or nucleus and *pi* is the proportion of a given splice variant among all detected isoforms in the cell or nucleus. This value is higher when isoforms are expressed at comparable levels than when some variants are disproportionately expressed. The value also increases with the number of isoforms detected.

### *In situ* hybridization

Methods were based on a prior publication.^24^ DNA templates were generated using forward primers with the SP6 promoter sequence at the 5′ end and reverse primers with the T7 promoter sequence at the 5′ end (Supplemental Table 2). Digoxigenin-labeled RNA probes were synthesized by SP6 RNA polymerase (#10810274001, Roche, Basel, Switzerland) or T7 RNA polymerase (#10881767001, Roche). Probes synthesized by T7 RNA polymerase coded the antisense sequence and thus were used for target RNA detection. Probes synthesized by SP6 RNA polymerase coded the sense sequence and thus were used for negative control experiments. Kidneys from PBS-perfused mice were fixed with 4% PFA and sectioned at 15 μm thickness. Sections were incubated with 10 μg/mL proteinase K for 20 minutes, 0.375% acetic anhydride solution for 10 minutes, and 0.5 μg/mL RNA probes in hybridization buffer (50% formaldehyde, 5× saline sodium citrate buffer [S8015, Sigma-Aldrich, St. Louis, MO], 1% sodium dodecyl sulfate, 50 μg/mL yeast tRNA, 50 μg/mL heparin) at 68°C overnight. Sections were then incubated with anti-digoxigenin-AP antibody (1:4,000; #11093274910, Roche) at 4°C overnight and developed with BM Purple (#11442074001, Roche) by overnight incubation. Sections were fixed in 4% PFA once again for 30 minutes and covered with mounting medium and coverslips.

### Quantitative PCR

Complementary DNA was subjected to real-time quantitative PCR with iTaq Universal SYBR Green Supermix (#1725121, Bio-Rad, Hercules, CA) and CFX Connect Real-Time PCR Detection System (Bio-Rad). Primer sequences are described in Supplemental Table 3. Gene expression was normalized to β-actin mRNA levels.

### Statistical analysis

Unpaired two-tailed Welch’s t test was performed to compare the mean of two groups. The level of statistical significance was set at *p* < 0.05. Data are presented as mean ± SD to show the variation of the data and mean ± 1.96 SEM to show the uncertainty of the mean. Error bars in each figure are explained in each figure legend.

Additional methods are provided in the Supplemental Material.

## Results

### Single-cell and single-nucleus long-read RNA-seq of healthy mouse kidneys

Kidneys from three healthy C57BL/6J adult male mice were subjected to both single-cell and single-nucleus long-read RNA-seq (Figure 1A). The median read length was 757–884 bases (Figure 1B, Supplemental Table 1), and the number of reads was 119–168 million reads per sample (Supplemental Table 1). Compared to single-cell samples, single-nucleus samples yielded a higher proportion of intronic reads but a comparable number of exonic reads, because single-nucleus samples produced more total reads when each sample was sequenced on a single flow cell (Figure 1C). Full-length transcripts from single-cell samples were mapped to the reference genome at a higher rate than those from single-nucleus samples (Figure 1D), possibly because single-nucleus preparations are more likely to capture transcripts that are still undergoing RNA processing.^25^

**Figure 1.**
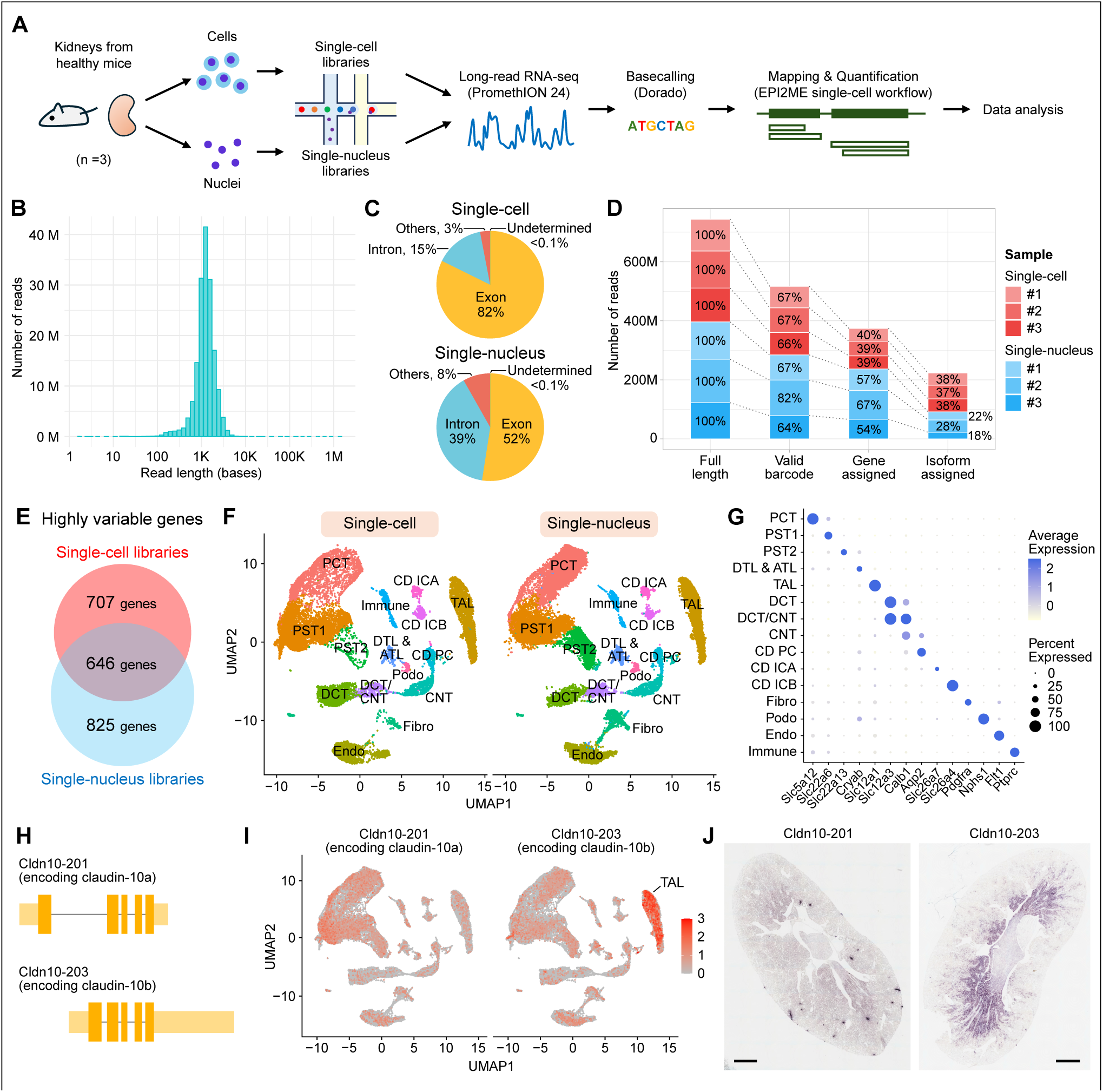
Single-cell and single-nucleus long-read RNA-seq of healthy mouse kidneys. (A) Experimental design. (B) Histogram of the read length of a representative sample. (C) Percentage of reads mapped to exons, introns and other regions. (D) Number of reads that were sequenced and mapped. Numbers shown in the bars represent the percentage of reads relative to the total number of full length reads in each dataset. (E) Number of highly variable genes that were unique to each dataset or common to both. (F) Uniform manifold approximation and projection (UMAP) plot of the single cells (left) and single nuclei (right) after integration and quality control. (G) Dot plot showing the expression pattern of cell type marker genes. (H) Schematic of two RNA isoforms of the gene *Cldn10*. Thick bars represent coding sequences, and thin bars represent untranslated regions. (I) UMAP presentation of the expression of Cldn10-201 and Cldn10-203 isoforms. (J) *In situ* hybridization of Cldn10-201 and Cldn10-203 isoforms on kidney sections from healthy mice. Scale bar, 1 mm. ATL, ascending thin limb of Henle’s loop; CD ICA, collecting duct – type A intercalated cell; CD ICB, collecting duct – type B intercalated cell; CD PC, collecting duct – principal cell; CNT, connecting tubule; DCT, distal convoluted tubule; DTL, descending thin limb of Henle’s loop; Endo, endothelial cell; Fibro, fibroblast; Immune, immune cell; K, kilo (thousand); M, million; PCT, proximal convoluted tubule; Podo, podocyte; PST, proximal straight tubule; TAL, thick ascending limb of Henle’s loop.

Single-cell and single-nucleus RNA-seq datasets differ systematically in gene expression profiles even within the same cell type, partly because of the abundance of mitochondrial transcripts in single-cell data. To integrate these datasets and cluster cells according to cell types, we identified highly variable genes that are common to both datasets (Figure 1E) and clustered cells based on the principal component analysis of the expression of these genes. Unsupervised clustering identified major kidney cell types after satisfactory integration of the single-cell and single-nucleus RNA-seq datasets, showing no method-dependent separations of clusters (Figure 1F). Cell type markers commonly used in single-cell and single-nucleus short-read RNA-seq were able to distinguish cell types in the long-read RNA-seq datasets as well (Figure 1G). Compared to single-cell samples, single-nucleus samples recovered a higher proportion of fibroblasts (5.2% in single-nucleus vs. 1.7% in single-cell samples) with only a modest reduction in the detection of immune cells (2.6% in single-nucleus vs. 3.4% in single-cell samples).

We next asked whether the datasets would recapitulate previously reported differences in the isoform usage among cells. The gene *Cldn10* encodes claudin-10, which is a tight junction component expressed as two isoforms generated by alternative splicing: claudin-10a and claudin-10b (Figure 1H).^9^ Claudin-10a forms anion-selective pores and is the major isoform in the kidney cortex, whereas claudin-10b creates cation-selective pores and is heavily expressed in the medulla and especially in the thick ascending limbs (TAL) of the loop of Henle.^9^ In our long-read dataset, isoform Cldn10-203 (encoding claudin-10b) was highly expressed in the TAL, which was in contrast to isoform Cldn10-201 (encoding claudin-10a) (Figure 1I). *In situ* hybridization confirmed the widespread expression of isoform Cldn10-201 in both the cortex and the medulla as well as the localization of isoform Cldn10-203 in the medulla of kidneys from healthy mice (Figure 1J).

Collectively, single-cell and single-nucleus long-read RNA-seq data from healthy mouse kidneys were successfully integrated and recapitulated known isoform usage patterns.

### Unique profiles of single-cell and single-nucleus long-read RNA-seq datasets

Next, we compared the single-cell and single-nucleus long-read RNA-seq datasets to explore transcriptomic differences arising from the methodology. At the gene level, most genes were detected in comparable proportions of cells and nuclei (Figure 2A). For 96.5% of all detected genes, the percentages of cells and nuclei expressing the gene differed by less than 20%. This observation is consistent with our previous comparison of single-cell and single-nucleus short-read RNA-seq in healthy kidneys.^26^ However, 37% of all genes showed significantly higher expression in single nuclei than single cells (Figure 2B), suggesting greater detection sensitivity with single-nucleus preparation. Genes enriched in single-cell samples included those related to mitochondria, ribosome and heat shock response (Figure 2C), which reflect the capture of cytoplasmic organelles and artifactual transcriptional stress responses induced by single-cell dissociation. The latter was missing from the single-nucleus dataset because nuclear suspensions are derived from snap frozen kidney samples.

**Figure 2.**
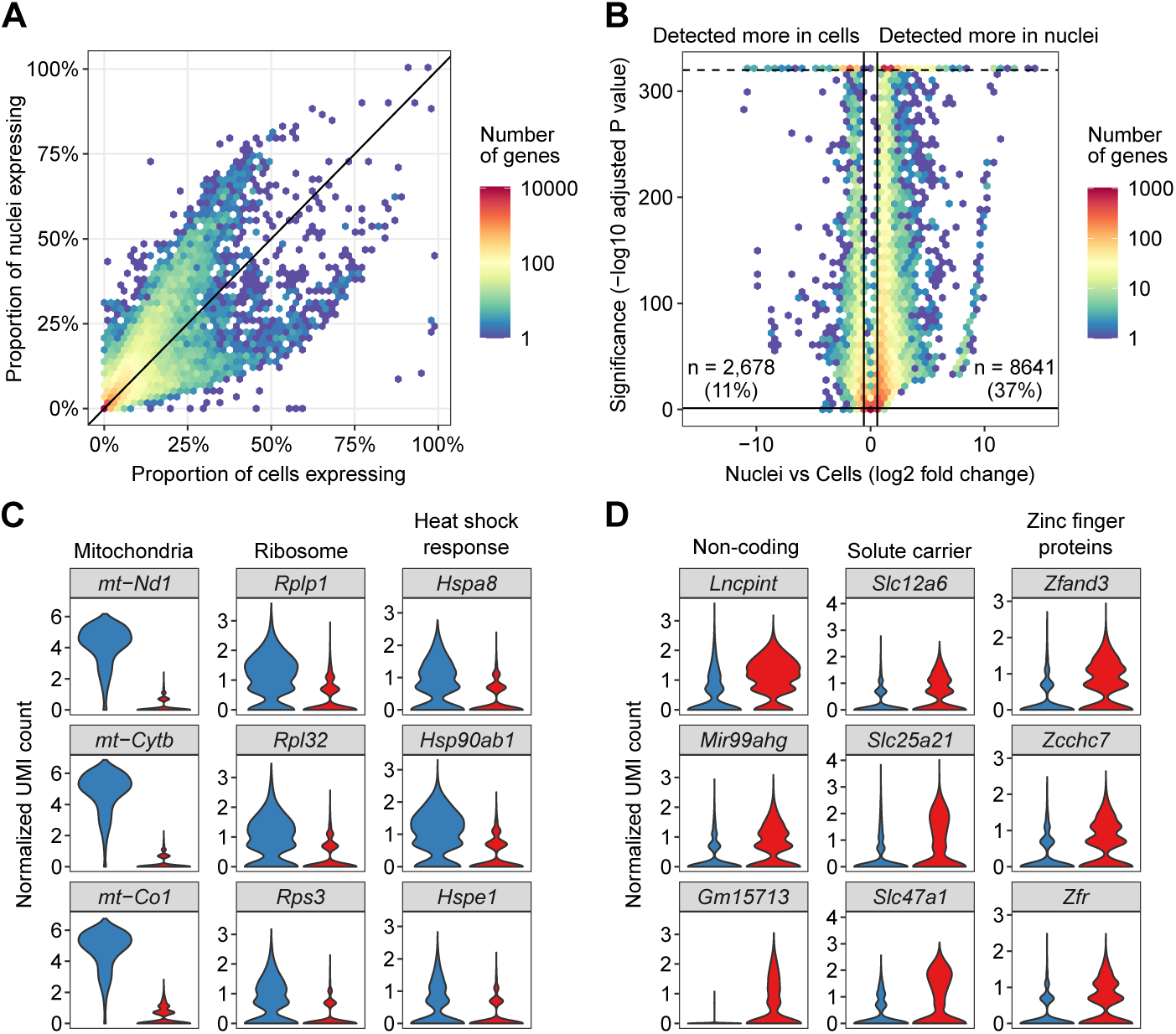
Gene expression profiles of single-cell and single-nucleus long-read RNA-seq datasets. (A) Binned scatter plot showing the proportion of cells (x axis) or nuclei (y axis) that express each gene. (B) Binned scatter plot of differentially detected genes showing that 11% and 37% of all detected genes were detected at a significantly higher rate in cells and nuclei, respectively. Bold lines indicate the thresholds for significance determination (fold change, ±1.5; adjusted p value, 0.05). The dashed line represents the smallest calculable adjusted p value. (C) Violin plot showing the expression of representative genes enriched in cells. (D) Violin plot showing the expression of representative genes enriched in nuclei.

Genes enriched in single-nucleus samples included cell-identity determining solute carriers and transcription factors, which was also consistent with our previous study (Figure 2D).^26^

At the isoform level, single-cell and single-nucleus long-read RNA-seq captured different patterns of alternatively spliced isoforms, as exemplified by isoforms of cell type marker genes (Figure 3A).

**Figure 3.**
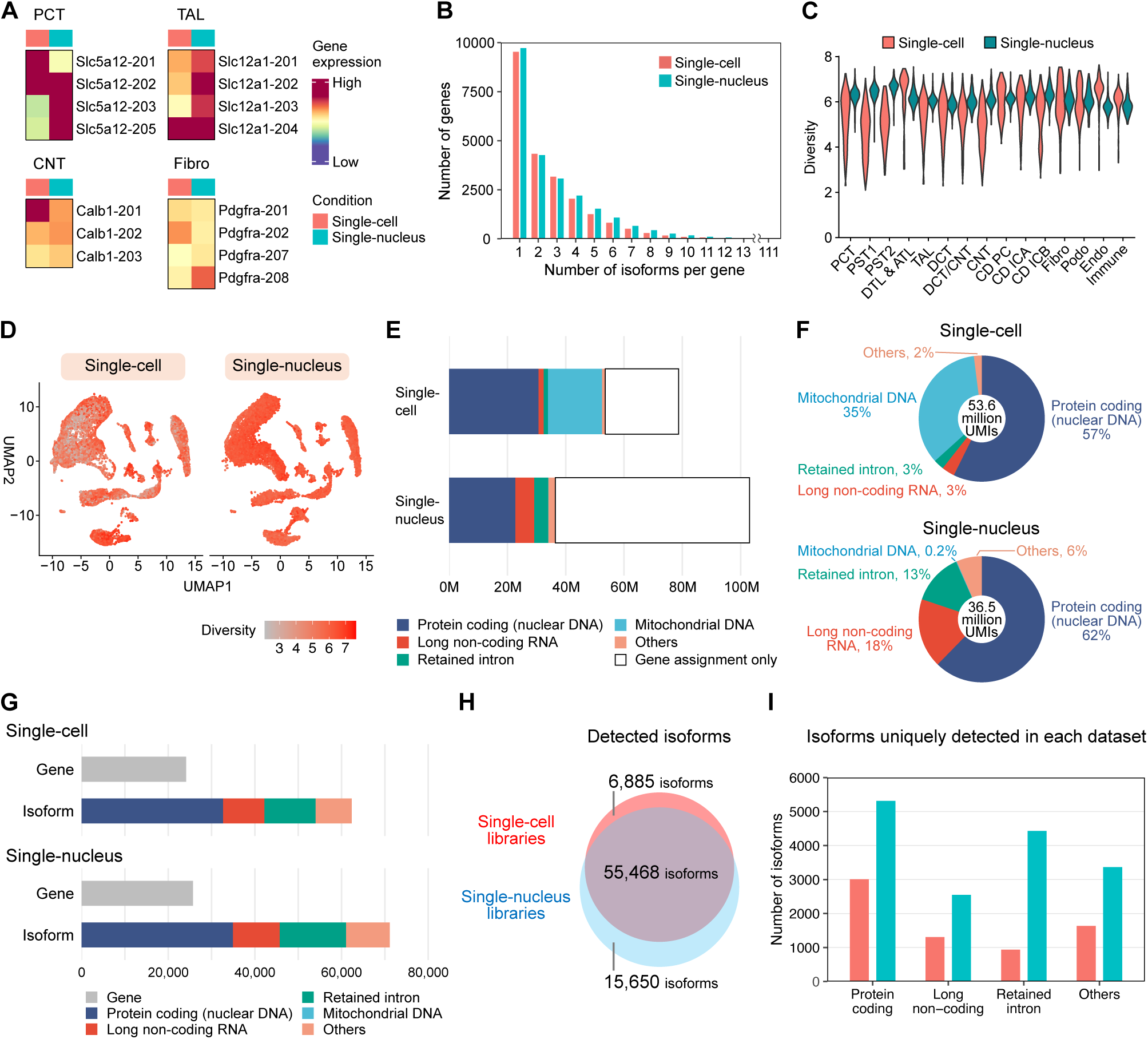
Unique isoform profiles of single-cell and single-nucleus long-read RNA-seq datasets. (A) Expression heatmap of cell type marker gene isoforms. (B) Number of isoforms detected per gene. (C, D) Isoform expression diversity per cell and nucleus. (E) Unique molecular identifier (UMI) counts and their types. (F) Distribution of isoform types among transcripts with isoform assignment. (G) Number of detected genes and isoforms. (H) Venn diagram of detected isoforms in single-cell and single-nucleus libraries. (I) Types of isoforms uniquely detected in each dataset.

Nuclear RNA processing, including post-transcriptional splicing, may account for the different isoform composition between cells and nuclei.^27^ The number of detected splice variants per gene was less than 9 in 95% of all detected genes in both datasets, with 111 variants for *Gas5* being the maximum (Figure 3B). The distribution was comparable between both datasets (Figure 3B). However, isoform expression diversity per cell or nucleus, defined by Equation 1 (see Methods), was consistently high across all cell types only in single nuclei, whereas it varied more among individual cells in most cell types in single cells (Figure 3C–D).

To determine whether the consistently high level of isoform expression diversity in single nuclei derived simply from a larger sequencing output, we analyzed the RNA molecule counts (technically, unique molecular identifier [UMI] counts) and their types in these datasets post-integration and quality control. The single-nucleus dataset contained 31% more UMIs (103.1 million vs. 78.7 million UMIs) but 32% fewer UMIs with isoform identification than the single-cell dataset (36.5 million vs. 53.6 million UMIs, Figure 3E). In the single-cell dataset, more than one third of the UMIs with isoform identification were the transcripts of merely 13 mitochondrial genes with no splice variants (Figure 3F), leaving a similar number of non-mitochondrial gene transcripts compared to that in the single-nucleus dataset. Despite the comparable non-mitochondrial UMI count in both datasets, the single-nucleus dataset contained 14% more splice variants than the single-cell dataset (71,118 vs. 62,353 splice variants, Figure 3G). Protein-coding isoforms constituted the largest category among the 15,650 and 6,885 isoforms that were uniquely detected in the single-nucleus and single-cell datasets, respectively (Figure 3H–I). The single-nucleus dataset contained more unique isoforms than the single-cell dataset in all three major categories: protein-coding isoforms, long non-coding isoforms and isoforms with retained introns (Figure 3I).

Together, single nuclei exhibited a consistently high level of isoform expression diversity across all cell types as compared to single cells not because of a larger sequencing output but likely driven in part by the detection of a wider variety of isoforms the scarcity of mitochondrial gene transcripts and detection of nascent unprocessed nuclear mRNAs.

### Single-nucleus long-read RNA-seq of kidneys from healthy and post-IRI mice

We next used single-nucleus long-read RNA-seq to investigate how acute kidney injury affects alternative splicing in the kidney. Single-nucleus long-read RNA-seq data from kidneys collected 2 days after bilateral IRI were compared with those from healthy kidneys (Figure 4A). After successful integration of the two datasets, unsupervised clustering based on gene expression identified 18 major cell types (Figure 4B). Injured proximal tubule cells and proliferating proximal tubule cells were found primarily in post-IRI kidneys (Figure 4C–D). We also clustered nuclei based on isoform expression alone (Figure 4E–F). Cell type annotations were mostly consistent between gene-based and isoform-based clustering (Figure 4G), but gene-based clustering provided clearer separation between clusters, possibly because of the higher number of UMIs per nucleus in the gene expression count matrix (Figure 1D). Therefore, gene-based clustering was used for downstream analysis.

**Figure 4.**
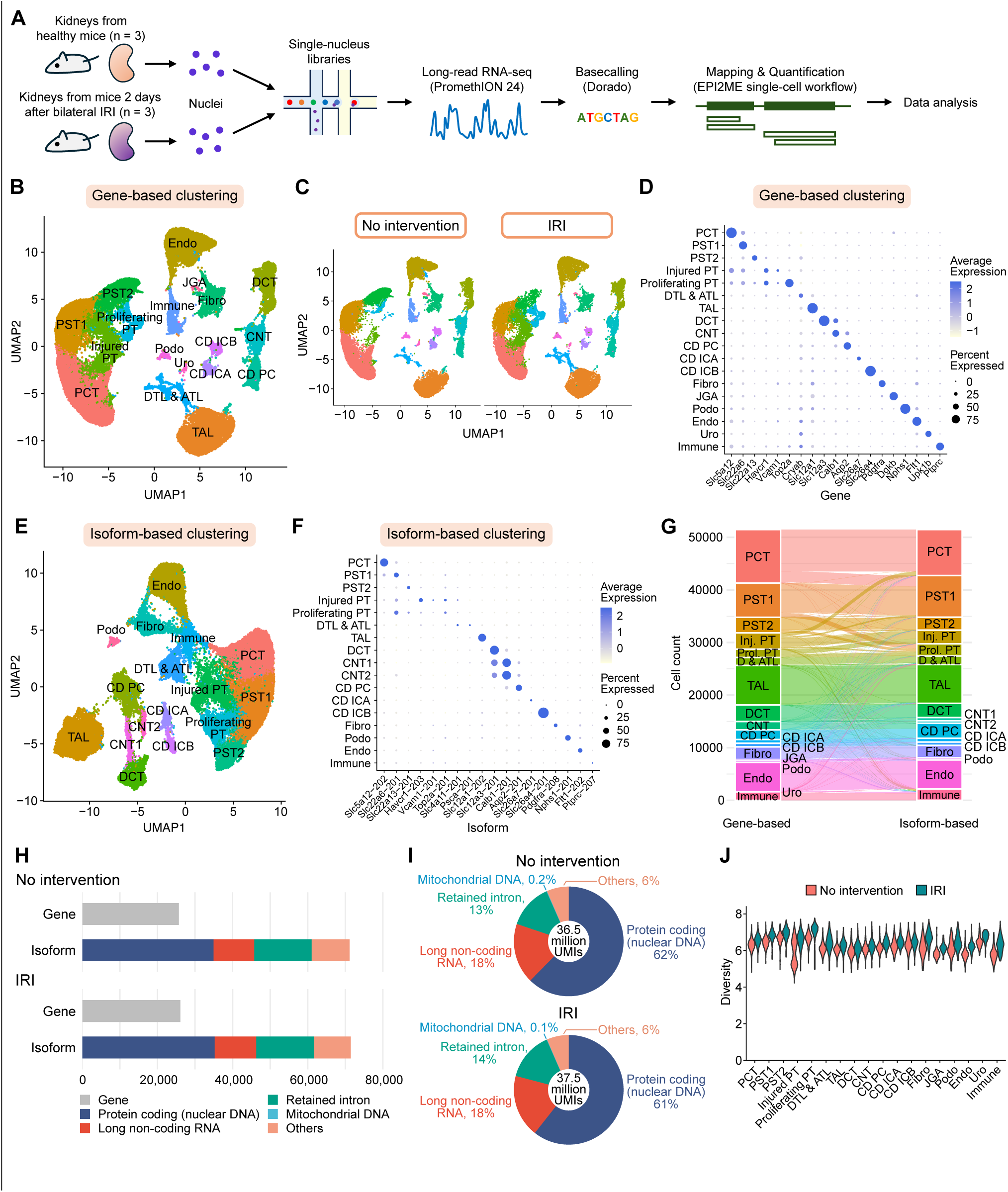
Single-nucleus long-read RNA-seq of kidneys from healthy and post-IRI mice. (A) Experimental design. (B) Uniform manifold approximation and projection (UMAP) plot of the integrated dataset clustered based on the gene expression. (C) UMAP plots of nuclei from control kidneys (left) and kidneys following IRI (right). (D) Dot plot showing the expression pattern of cell type marker genes. (E) UMAP plot of the integrated dataset clustered based on the isoform expression. (F) Dot plot showing the expression pattern of cell type marker isoforms. (G) Alignment between gene-based and isoform-based clusters. (H) Number of detected genes and isoforms. (I) Distribution of isoform types among transcripts with isoform assignment. (J) Isoform expression diversity per nucleus. D & ATL, descending and ascending thin limbs of Henle’s loop; inj., injured; IRI, ischemia-reperfusion injury; JGA, juxtaglomerular apparatus; prol., proliferating; PT, proximal tubule; Uro, uroepithelial cell.

We first assessed overall isoform expression metrics in healthy kidneys and post-IRI kidneys. The number of isoforms detected in the two sample groups were comparable (Figure 4H), and the isoform type composition was similar (Figure 4I). However, kidneys following IRI exhibited greater isoform expression diversity per nucleus (Equation 1) across all cell types (Figure 4J), reflecting the IRI-induced changes in isoform regulation in kidneys.

### Isoform-level regulation of gene expression in proximal tubule cells

Gene expression changes in proximal tubule cells after IRI have been studied extensively, but isoform regulation in these cells after IRI remain poorly characterized. We analyzed whether IRI induces altered isoform regulation in proximal tubule cells (Figure 5A). Using DTUrtle,^22^ we identified 2,414 genes with differential isoform usage, meaning a statistically significant change in the proportion of isoforms expressed in the nuclei (Figure 5B). More than half of these genes had exactly two detected isoforms, and the change in the proportion of two isoforms was statistically significant in all genes (Figure 5B). Among these genes with two isoforms, 449 genes (32%) showed a shift in isoform usage from a protein-coding isoform to another protein-coding isoform, 433 genes (31%) from a protein-coding isoform to a non-protein-coding isoform, 409 genes (29%) from a non-protein-coding isoform to a protein-coding isoform, and 117 genes (8%) from a non-protein-coding isoform to another non-protein-coding isoform (Figure 5C).

**Figure 5.**
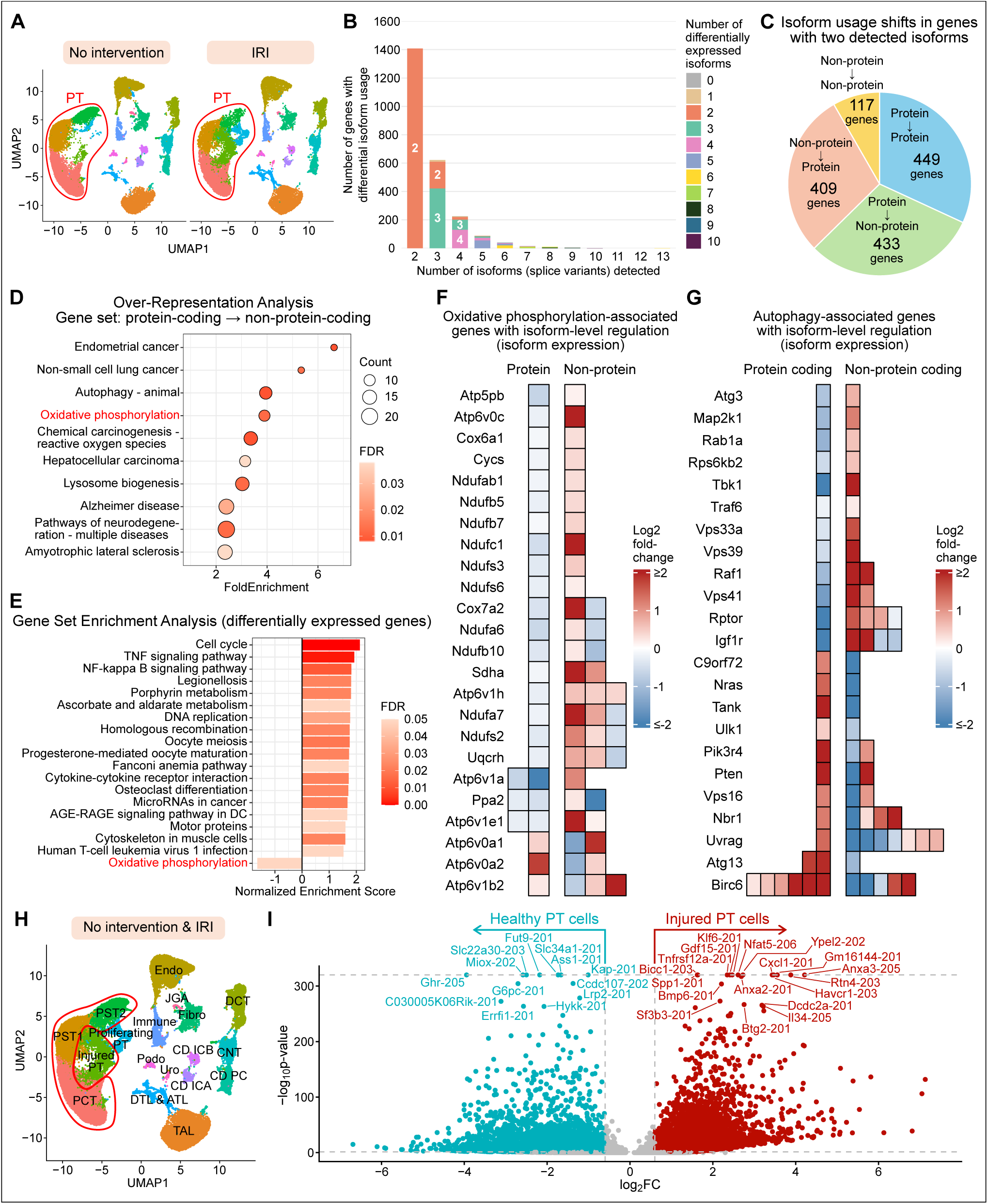
Isoform-level regulation of gene expression in proximal tubule cells. (A) Two groups of proximal tubule cells (red circles) compared for isoform usage. (B) Distribution of the numbers of detected isoforms and differentially expressed isoforms among genes with differential isoform usage. (C) Distribution of isoform usage shifts in genes with two detected isoforms. (D) KEGG gene categories identified by over-representation analysis of genes with a shift from a protein-coding isoform to a non-protein-coding isoform. FDR, false discovery rate. (E) KEGG gene categories identified by gene set enrichment analysis for transcriptomic difference between proximal tubule cells in healthy kidneys and those in kidneys following IRI. DC, diabetic complications. (F) Isoform expression heatmap of oxidative phosphorylation–associated genes with isoform-level regulation. (G) Isoform expression heatmap of autophagy-associated genes with isoform-level regulation. (H) Two groups of cells that were analyzed for differentially expressed isoforms (red circles). (I) Differentially expressed isoforms. Blue and red colors indicate statistically higher isoform expression in healthy and injured proximal tubule (PT) cells, respectively. AGE, advanced glycation end-products; FC, fold change; FDR, false discovery rate; RAGE, receptor for advanced glycation end-products; TNF, tumor necrosis factor.

To explore the biological implications of these changes in proximal tubule cells after IRI, we performed over-representation analysis on these sets of genes. Several KEGG gene categories were significantly enriched among genes that shifted from a protein-coding isoform to a non-protein-coding isoform (Figure 5D) or to another protein-coding isoform (Supplemental Figure 3). In addition to this isoform-level analysis, we also performed gene set enrichment analysis on gene-level expression differences between proximal tubule cells from healthy and post-IRI kidneys (Figure 5E). Notably, oxidative phosphorylation was downregulated at both the gene level (Figure 5E) and the isoform level, the latter through shifts from protein-coding to non-protein-coding isoforms (Figure 5D).

We further investigated IRI-induced isoform-level expression changes in oxidative phosphorylation– associated genes. Among 141 oxidative phosphorylation–associated genes defined in KEGG, 89 genes were detected in our samples. Among these, 24 genes showed isoform-level regulation, defined as concordant up- or down-regulation of all protein-coding isoforms accompanied by opposite regulation of at least one non-protein-coding isoform (Figure 5F). Notably, 21 of these 24 genes (88%) showed downregulation of all protein-coding isoforms, indicating that oxidative phosphorylation-associated genes were downregulated at the isoform level as well. This shift from protein-coding toward non-protein-coding isoforms did not reflect a global response to IRI, as the total expression of protein-coding and non-protein-coding isoforms was comparable between healthy and post-IRI kidneys (Figure 4I). Moreover, this directional bias was not common to all gene categories listed in Figure 5D, because some categories contained comparable numbers of genes with upregulated and downregulated protein-coding isoforms, as exemplified by autophagy-associated genes (Figure 5G).

In addition, we compared the use of splice variants in injured proximal tubule cells and healthy proximal tubule cells (Figure 5H). While isoform usage shifts were less obvious than in the analysis above, differentially expressed isoforms were identified, recapitulating the established differentially expressed genes at isoform resolution (Figure 5I). This observation suggests that gene-level regulation, such as the expression of a completely different gene set, could be playing a larger role than isoform-level regulation of expression in injured proximal tubule cells. Taken together, proximal tubule cells exhibited isoform-level regulation of gene expression following IRI, as exemplified by genes associated with oxidative phosphorylation.

### Isoform usage shifts of *Fut9* and *Prrc2b* genes after IRI

We sought to identify specific genes exhibiting prominent isoform usage shifts in proximal tubule cells following IRI. Among the 2,414 genes with differential isoform usage in proximal tubule cells (Figure 5B), *Fut9* showed a pronounced isoform usage shift, with decreased expression of the protein-coding isoform Fut9-201 and increased expression of the non-protein-coding isoform Fut9-203 (Figure 6A). *Fut9* codes for fucosyltransferase 9, an enzyme that is conserved across mammals and catalyzes the α1,3-fucosylation of N-acetylglucosamine residues on glycan chains, influencing diverse processes including immune cell recognition and cell adhesion.^28^ Quantitative PCR verified the decreased expression of the major protein-coding isoform Fut9-201 after IRI (Figure 6B). Western blot confirmed the decreased amount of fucosyltransferase 9 protein in kidneys following IRI (Figure 6C). Such change could not be inferred from the comparable *Fut9* gene expression, which includes all transcripts regardless of isoform assignment (Figure 6D–E), highlighting the power of single-cell and single-nucleus long-read RNA-seq in detecting such isoform usage shifts.

**Figure 6.**
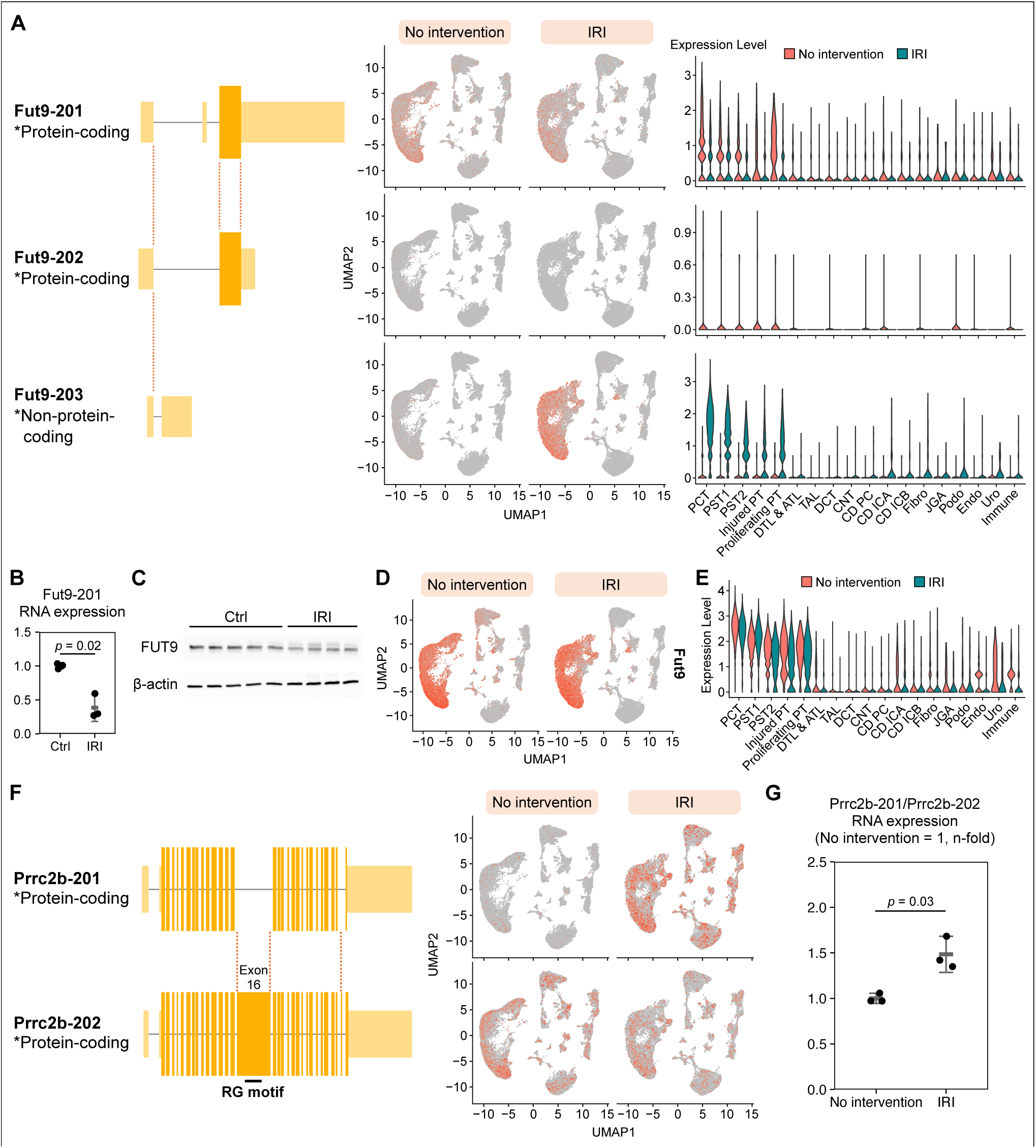
Isoform usage shifts of *Fut9* and *Prrc2b* genes after IRI. (A) Left; schematic of *Fut9* gene structure and regions transcribed into mRNA isoforms. Dark yellow indicates the protein-coding regions. Middle; feature plots showing the expression of each *Fut9* isoform. Right; violin plots showing the expression of each *Fut9* isoform, stratified by cell type. (B) Fut9-201 mRNA expression level (relative quantity) verified by quantitative PCR. (C) Western blot of fucosyltransferase 9 (FUT9) and β-actin in control kidneys and kidneys following IRI. (D) Feature plots showing the *Fut9* gene expression. (E) Violin plot showing the *Fut9* gene expression, stratified by cell type. (F) Left; schematic of *Prrc2b* gene structure and regions transcribed into mRNA isoforms. Dark yellow denotes the protein-coding regions. Right; feature plots showing the expression of each *Prrc2b* isoform. RG motif, arginine-glycine motif. (G) Prrc2b-201 to Prrc2b-202 mRNA expression level ratio verified by quantitative PCR.

Genes with isoform switches between two protein-coding isoforms with distinct functions were also detected. A notable example is *Prrc2b*, which encodes proline-rich coiled-coil 2B (PRRC2B).

PRRC2B is an RNA-binding protein that regulates gene expression posttranscriptionally through RNA splicing, transport, modification, translation and degradation.^29^ Of the two major alternatively spliced isoforms of *Prrc2b* mRNA, the shorter isoform lacks exon 16, which harbors arginine-glycine repeats (RG motif) that serve as the RNA binding site (Figure 6F, left), thereby limiting the shorter isoform’s regulatory role compared to the longer isoform.^30^ In our single-nucleus RNA-seq data, the longer isoform (Prrc2b-202) was dominant in healthy kidneys, whereas the shorter isoform (Prrc2b-201) was highly expressed in multiple cell types after IRI (Figure 6F, right). Quantitative PCR verified the increase in the ratio of Prrc2b-201 to Prrc2b-202 RNA expression levels in kidneys after IRI (Figure 6G).

## Discussion

In this study, we investigated differences in isoform expression between healthy and acutely injured mouse kidneys by performing long-read RNA-seq at single-cell resolution. We found that genes associated with oxidative phosphorylation were regulated not only at the gene expression level but also through shifts in isoform composition in proximal tubule cells after IRI. Beyond oxidative phosphorylation–associated genes, we identified *Fut9* and *Prrc2b* as genes exhibiting prominent shifts in isoform usage after IRI.

Oxidative phosphorylation–associated genes are downregulated in proximal tubule cells in multiple kidney diseases.^5,7,31^ However, whether these genes are regulated at the isoform level was previously unknown. Our single-nucleus long-read RNA-seq approach identified shifts from protein-coding transcripts to non-protein-coding transcripts in proximal tubule cells after IRI across a set of oxidative phosphorylation–related genes, many of which encode subunits of mitochondrial Complex I–V. These shifts were concordant with the gene-level downregulation of oxidative phosphorylation– associated genes, suggesting an additional layer of regulation.

Although multiple studies have characterized alternative splicing of specific genes in the kidney,^8–10^ comprehensive analysis of the whole transcriptome at the isoform level had been challenging due to the limited read length of short-read RNA-seq. Long-read RNA-seq has enabled the identification of transcripts at the isoform level with substantially improved sequencing accuracy compared to earlier generations of the technology.^11^ In our study, the median Phred quality score of the reads was 21.9 or higher for the single-cell datasets and 24.1 or higher for the single-nucleus datasets (Supplemental Table 1), corresponding to median base-call accuracies of 99.4% and 99.6%, respectively. This accuracy is lower than that of conventional short-read sequencers but was sufficient for gene- and isoform-level read assignment. The median read length ranged from 757 to 884 bases (Figure 1B, Supplemental Table 1), which was substantially longer than short reads.

These lengths were comparable to those reported in previous single-cell long-read RNA-seq studies of other tissues^15^ and bulk long-read RNA-seq of mouse kidneys.^32^

This study also characterized the different profiles of single-cell and single-nucleus long-read datasets. Although single-nucleus datasets contained a higher proportion of intronic reads (Figure 1C), single-nucleus long-read RNA-seq detected more distinct isoforms (Figure 3G) and greater isoform expression diversity (Figure 3C) than single-cell long-read RNA-seq. This may reflect the abundance of mitochondrial DNA–derived reads in single-cell datasets as well as ongoing RNA processing in the nucleus. One might expect single-cell long-read RNA-seq to capture a higher proportion of mature mRNAs and thus to yield a transcriptome that more closely corresponds to protein expression. However, single-cell data are susceptible to artifactual transcriptional changes induced by the single cell isolation process (Figure 2C). Meanwhile, the non-coding isoforms captured in single-nucleus datasets may be useful for studying alternative splicing and other forms of transcriptional regulation.^33^ The choice between the two approaches should therefore depend on the research question.

This study has several limitations. First, the biological consequences of the *Fut9* and *Prrc2b* isoform usage shifts were not investigated in this study. Fucosyltransferase 9 modulates immune cell recognition,^28^ while PRRC2B regulates cell cycle and cell development,^29,34^ among other functions. Whether and how these isoform usage shifts contribute functionally in acutely injured kidneys warrants further investigation. Second, only male mice were used for the experiments because they are known to be more susceptible to renal IRI.^35^ Finally, sequence reads were mapped to transcripts in the GENCODE/Ensembl reference annotation, so novel isoforms absent from the reference annotation were not captured in our analysis. Including these isoforms in the analysis could increase the number of reads assigned to isoforms (Figure 1D).

In summary, single-nucleus long-read RNA sequencing revealed injury-induced shifts in isoform usage in kidney cells, adding a layer of transcriptomic information that is largely inaccessible to conventional gene-level short-read analysis.

## Declarative Statements

All animal experiments were conducted in accordance with the NIH Guide for the Care and Use of Laboratory Animals. All protocols were approved by the Institutional Animal Care and Use Committee at Washington University in St. Louis (protocol #23-0460).

## Use of Artificial Intelligence

Claude Opus 5 (Anthropic) was used to assist with coding for data analysis and figure generation, with revising manuscript text, and with checking the manuscript for completeness, errors and inconsistencies. The authors reviewed and verified all such output and are fully responsible for the content.

