## Supplemental for "Single-nucleus long-read RNA sequencing reveals isoform usage shifts in mouse acute kidney injury"

### Supplemental Material

### Single-nucleus long-read RNA sequencing unravels isoform-level regulation of gene expression in normal and acutely injured kidneys

**Yasuhiro Oda, Megan L. Noonan, Eryn E. Dixon, Haojia Wu, and Benjamin D. Humphreys**

Table of contents

- Supplemental Methods.
- Supplemental Table 1. Key metrics of single-cell/nucleus long-read RNA sequencing data.
- Supplemental Table 2. Primer sequence for quantitative PCR.
- Supplemental Figure 1. Barcode rank plots for single-cell and single-nucleus long-read RNA sequencing datasets before empty droplet removal.
- Supplemental Figure 2. Quality control of clusters in the integrated dataset of single-cell and single-nucleus long-read RNA sequencing data.
- Supplemental Figure 3. KEGG gene categories identified by over-representation analysis of genes with differential usage of two detected protein-coding isoforms.

**Supplemental Methods**

**Single cell isolation**

Single cells were isolated as described in the literature (Ide et al., 2022) with some modifications. Mice were euthanized and perfused with ice-cold PBS. Kidneys were collected and minced into small pieces with a razor blade. Samples were incubated in a digestion buffer that contains 0.3 mg/mL Liberase TM (#291963, Roche, Basel, Switzerland), 20 μg/mL DNase I (#11284932001, Roche) and 10 μg/mL hyaluronidase (H4272, Sigma-Aldrich, St. Louis, MO) at 37°C for 20 minutes. After 2 minutes incubation with bed blood cell lysis buffer, cells were incubated with trypsin buffer for 10 minutes followed by trypsin inactivation with 10% fetal bovine serum in PBS. Cells were mechanically dissociated using a syringe and a 20 G needle and filtered through 70 μm and 30 μm cell strainers. After checking the cell count and viability, cell suspensions were adjusted to 1,000 cells/μL with PBS supplemented with 0.04% bovine serum albumin.

**Single nucleus isolation**

Single nuclei were isolated as previously described (Kirita et al., 2020) with some modifications. Kidneys were snap-frozen after euthanasia and stored in −80°C. Kidneys were minced into small pieces with a razor blade and homogenized with a Dounce homogenizer (#885302-0002, DWK Life Sciences, Wertheim, Germany) in ice-cold Nuclei EZ Lysis buffer (NUC-101, Sigma-Aldrich) supplemented with protease inhibitor (#5892791001, Roche) and RNase inhibitor (N2615, Promega, Madison, WI; AM2696, Thermo Fisher Scientific, Waltham, MA). After 5-minute incubation on ice, the homogenate was filtered through a 200-μm cell strainer (#43-50200, pluriSelect, Leipzig, Germany), homogenized again and incubated with the buffer on ice for 5 minutes. The homogenate was passed through a 40-μm cell strainer (#43-50040, pluriSelect), pelleted by centrifugation, resuspended in the buffer and incubated on ice for 5 minutes. The cell suspension was centrifuged, pelleted, resuspended with PBS supplemented with 0.1% RNase inhibitor, and filtered through a 5-μm cell strainer (#43-50005, pluriSelect). After checking the nuclei count, nuclei suspensions were adjusted to 1,000 cells/μL.

**Western blot**

Kidneys were homogenized in radioimmunoprecipitation assay buffer with Protease/Phosphatase Inhibitor Cocktail (#5872, Cell Signaling Technology, Danvers, MA) using tubes containing zirconium-silicate beads (Lysing Matrix D; #116913100, MP Biochemicals, Santa Ana, CA) and a microtube homogenizer (BeadBug 6; D1036, Benchmark Scientific, Sayreville, NJ). After 15 minutes of incubation on ice and 10 minutes of centrifugation, supernatant was collected and stored at −80°C. Protein levels were quantified by Pierce BCA Protein Assay Kit (#23252, Thermo Fisher Scientific). Protein solution was mixed with sample buffer which contained Laemmli sample buffer (#1610747, Bio-Rad, Hercules, CA) and 2-mercaptoethanol followed by boiling at 95°C for 5 minutes. Proteins were separated with 4–20% polyacrylamide gel electrophoresis and transferred to polyvinylidene difluoride membranes (Immobilon-P PVDF Membrane; #IPVH00010, Millipore, Burlington, MA) in a buffer tank. To prevent nonspecific protein binding, membranes were shaken in PBS with 5% bovine serum albumin at room temperature for 1 hour. Membranes were incubated with primary antibodies in PBS for 1 h at room temperature, washed with Tris-buffered saline with 0.05% Tween 20 (TBS-T) three times for 5 minutes each, incubated with secondary antibodies in PBS for 1 hour at room temperature, and washed with TBS-T three times for 5 minutes each. Blots were visualized by Amersham ECL Prime Western Blotting Detection Reagent (RPN2236, Cytiva, Marlborough, MA) and ChemiDoc (Bio-Rad). Antibodies used in this study are as follows: mouse anti-fucosyltransferase 9 monoclonal antibody (1:1,000; #60230-1-Ig, Proteintech, Rosemont, IL), mouse anti-β-actin monoclonal antibody (1:1,000; #3700, Cell Signaling Technology), and goat anti-mouse immunoglobulins/horseradish Peroxidase (1:5,000; P0447, Dako, Glostrup, Denmark).

**Supplemental Table 1. Key metrics of single-cell/nucleus long-read RNA sequencing data.**

| Sample | Number of reads | Median read length (base) | Median read quality | UMI counts, min cutoff | UMI counts, max cutoff | Gene counts, max cutoff | Number of cells/nuclei |
|---|---|---|---|---|---|---|---|
| Single-cell, healthy | | | | | | | |
| #1 | 118,698,553 | 883 | 21.9 | 281 | 4,737 | 17,346 | 8,421 |
| #2 | 136,843,275 | 850 | 22.5 | 193 | 5,345 | 20,389 | 7,489 |
| #3 | 125,406,104 | 884 | 22.3 | 231 | 4,178 | 13,346 | 10,347 |
| Single-nucleus, healthy | | | | | | | |
| #1 | 143,822,610 | 811 | 24.9 | 514 | 4,645 | 13,539 | 11,130 |
| #2 | 168,052,329 | 794 | 24.2 | 538 | 4,495 | 12,597 | 12,682 |
| #3 | 145,004,964 | 765 | 24.4 | 367 | 4,552 | 12,820 | 12,853 |
| Single-nucleus, IRI | | | | | | | |
| #1 | 124,561,437 | 882 | 24.9 | 631 | 5,521 | 16,916 | 10,009 |
| #2 | 200,439,015 | 757 | 24.1 | 678 | 4,113 | 12,639 | 9,119 |
| #3 | 156,199,001 | 837 | 24.7 | 699 | 4,638 | 15,061 | 9,823 |

*Read quality: Phred quality score.
*UMI counts, minimum cutoff: UMI counts at the bottom of the cliff in barcode rank plots.
*UMI counts, maximum cutoff: upper 1 percentile of the remaining cells.
*Gene counts, maximum cutoff: upper 1 percentile of the remaining cells.
*Number of cells/nuclei represents those remained after quality control.

**Supplemental Table 2. Primer sequence for creating probe templates for *in situ* hybridization.**

| Isoform | Forward | Reverse |
|---|---|---|
| Cldn10-201 | CATTTAGGTGACACTATAGATGGGCTGGCTT TAACATTCG | TAATACGACTCACTATAGGGGATGAAGAGAGCAGC CCCTAA |
| Cldn10-203 | CATTTAGGTGACACTATAGCTTGGAAATCGTC GCCTTCG | TAATACGACTCACTATAGGGCAGACGTGGGTCCGT TGTAT |

**Supplemental Table 2. Primer sequence for quantitative PCR.**

| Gene/isoform | Forward | Reverse |
|---|---|---|
| *Actb* | GAGGTATCCTGACCCTGAAGTA | GCTCGAAGTCTAGAGCAACATAG |
| Fut9-201 | CGCTGGGATCGCTTTAGAATG | AATGGGCGAAGAATGCCTTTG |
| Prrc2b-201 | GAAAGTGACCGCCCTCGAC | GGCAAGGCCTCATCTGGAG |
| Prrc2b-202 | TTGGGGAGAAGCCTGTTAGG | GGCAAGGCCTCATCTGGAG |

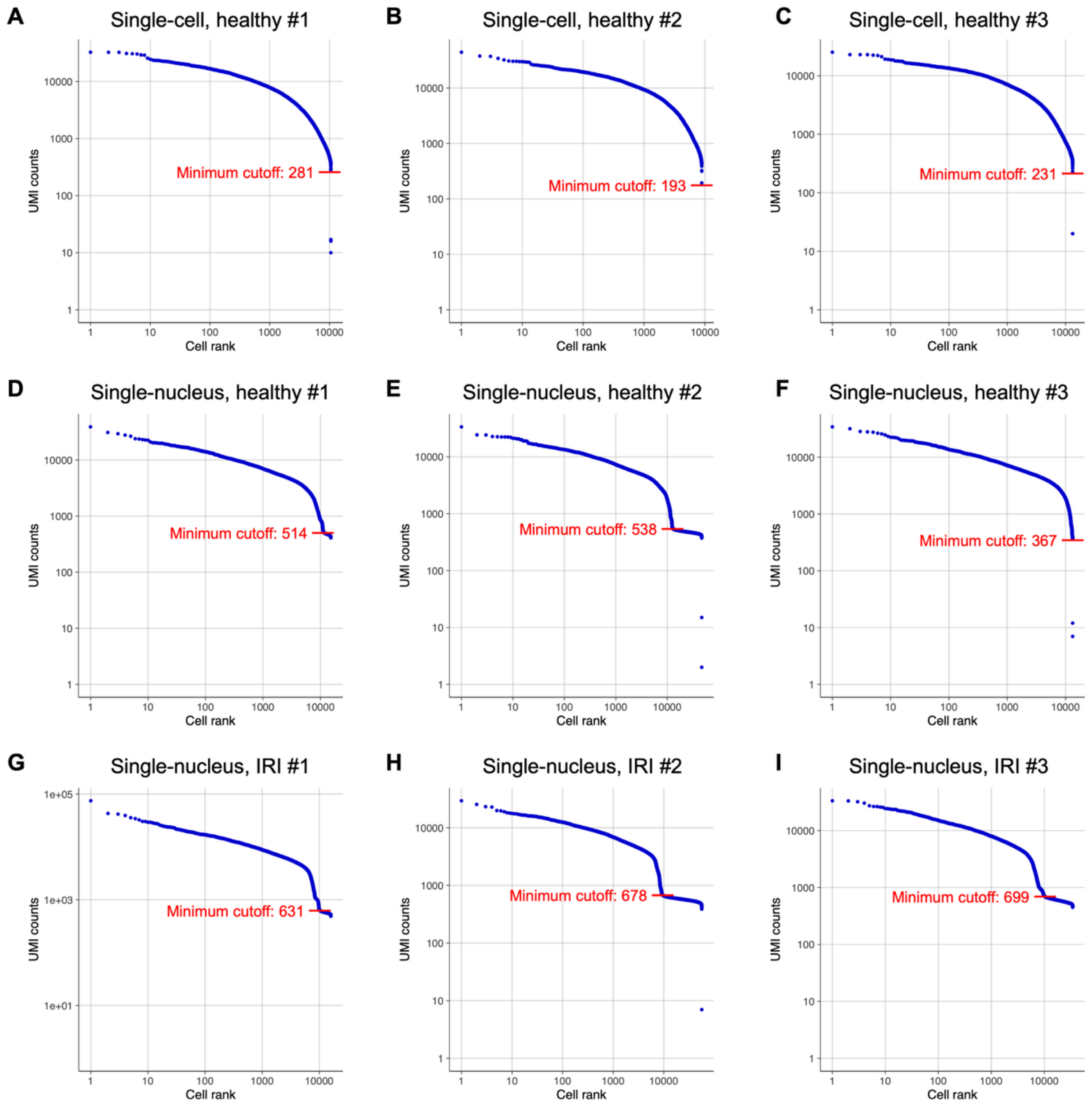


**Supplemental Figure 1. Barcode rank plots for single-cell and single-nucleus long-read RNA sequencing datasets before empty droplet removal.**
(A) Single-cell, healthy #1, (B) single-cell, healthy #2, (C) single-cell, healthy #3, (D) single-nucleus, healthy #1, (E) single-nucleus, healthy #2, (F) single-nucleus, healthy #3, (G) single-nucleus, IRI #1, (H) single-nucleus, IRI #2, (I) single-nucleus, IRI #3. Red bars show the unique molecular identifier (UMI) counts at the bottom of the cliff and were used as the minimum cutoff threshold.

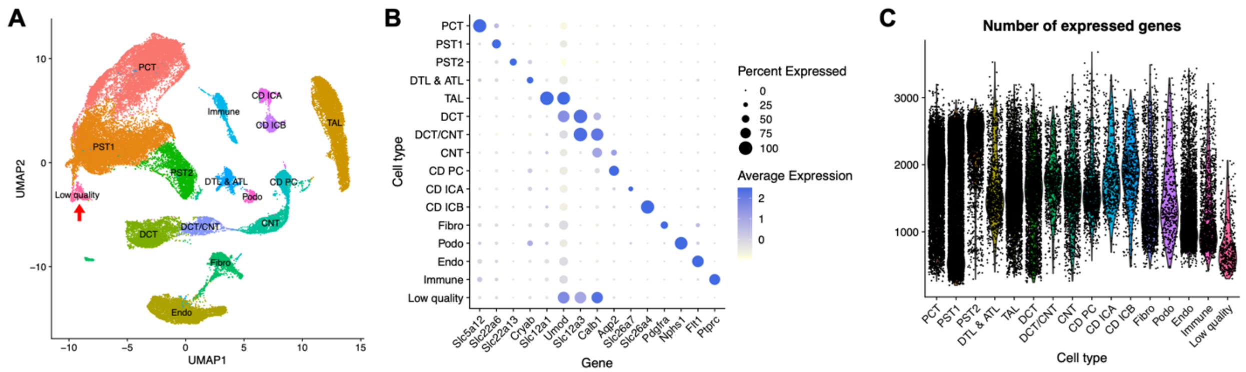


**Supplemental Figure 2. Quality control of clusters in the integrated dataset of single-cell and single-nucleus long-read RNA sequencing data.**

(A) Uniform manifold approximation and projection (UMAP) plot of the integrated dataset. Red arrow denotes the cluster of low-quality cells. (B) Dot plot showing the expression pattern of cell type marker genes in each cluster. (C) Number of expressed genes per cell, stratified by cell type.

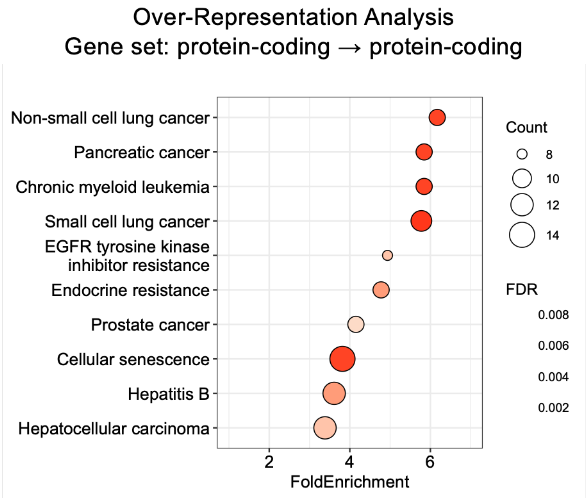


**Supplemental Figure 3. KEGG gene categories identified by over-representation analysis of genes with differential usage of two protein-coding isoforms.**
